# Lymph-node transcriptomics define prognostic immune states in mucosal melanoma and reveal IBA1 as a practical biomarker for improved prognosis

**DOI:** 10.1101/2025.09.20.677500

**Authors:** Angus Lane, Laura Blackwood, Kevin Donnelly, Yuting Lu, Alison Meynert, Jorge del Pozo, Gerry Polton, Laura Selmic, Jean-Benoit Tanis, David Killick, Joanna Morris, Inge Breathnach, Stefano Zago, Sara Gould, Scott Maxwell, Mickey Tivers, Davide Malucelli, Ana Marques, Katarzyna Purzycka, Matteo Cantatore, E. Elizabeth Patton, Kelly L. Bowlt Blacklock

**Affiliations:** Royal (Dick) School of Veterinary Studies and the Roslin Institute, Edinburgh, EH25 9RG, UK; MRC Human Genetics Unit, Institute of Genetics and Cancer, University of Edinburgh, Edinburgh EH4 2XU, UK; CRUK Scotland Centre, Institute of Genetics and Cancer, University of Edinburgh, Edinburgh EH4 2XU, UK; North Downs Specialist Referrals, 3&4 The Brewerstreet Dairy Business Park, Brewer Street, Bletchingley, RH1 4QP, UK; Department of Veterinary Clinical Sciences, The Ohio State University, 601 Vernon L. Tharp St., Columbus, OH 43210, USA; University of Liverpool. Leahurst Campus, Chester High Road, Neston, Wirral, CH64 7TE, UK; School of Biodiversity, One Health and Veterinary Medicine. University of Glasgow, Bearsden Road, Glasgow G61 1QH, UK; The Ralph Veterinary Referral Centre, Fourth Avenue, Globe Business Park, Marlow, SL7 1YG, UK; University of Bristol, Langford, Bristol, BS40 5DU, UK; Paragon Veterinary Referrals, Red Hall Crescent, Wakefield, WF1 2DF, UK; VetsNow, 123-145 North Street, Glasgow, G3 7DA, UK; Anderson Moores Veterinary Specialists, The Granary, Bunstead Barns, Hampshire, SO21 2LL, UK

**Keywords:** Mucosal melanoma, Canine, Oronasal, Transcriptome, Immunity, Metastasis, Lymph node

## Abstract

Mucosal melanoma (MM) is a rare and aggressive cancer in humans with poor prognosis and limited response to immunotherapy or targeted therapy. Progress has been hindered by the lack of immune-competent, translational models. Dogs develop oral mucosal melanoma (OMM), a biologically equivalent disease, making them a valuable companion animal model that shares the human environment and immune context.

In this study, we present the first transcriptomic analysis of regional lymph nodes in dogs with OMM, revealing that lymph nodes stratify into two distinct subgroups, independent of histopathological metastatic status at time of surgery, and which can be succinctly captured with a 35-gene signature. Notably, this stratification correlates with survival outcomes, providing unique early prognostic insights at the time of diagnosis.

Furthermore, we explore the immune landscape of these subgroups and identify IBA1+ monocyte/macrophage infiltration as a key distinguishing biomarker. Using immunohistochemistry (IHC) and digital pathology, we demonstrate that higher IBA1 expression is associated with the transcriptomic subgroups associated with improved survival. These findings highlight IBA1 as a potential biomarker for risk stratification, offering a clinically relevant tool for refining prognosis and guiding treatment decisions in canine OMM.

**Highlights:**

- **Transcriptomic profiling of canine OMM lymph nodes identifies two prognostic subgroups**, independent of histopathological metastatic status.
- **Lymph node transcriptomic stratification correlates with survival**, providing prognostic insights beyond conventional histopathology at the time of diagnosis.
- **IBA1+ monocyte/macrophage infiltration** is associated with improved survival, and shows strong diagnostic potential via immunohistochemistry, supporting its use as a clinically feasible stratification tool.

## Introduction

Mucosal melanoma (MM) is a rare and aggressive form of melanoma arising from melanocytes in sun-protected mucosal surfaces [1]. Unlike cutaneous melanoma, MM lacks well-defined precursor lesions, oncogenic driver mutations or effective therapeutics [2–6].

The dog is an ideal immunocompetent model for human MM because of the commonality of clinical, pathological, environmental, and genomic features [6–8]. Oral MM (OMM) is the most common oral tumour in dogs and, similar to human OMM, is characterised by local invasion, frequent recurrence after radical surgical resection, high metastatic propensity, rapid progression from localised to advanced-stage disease, and limited response to current therapeutic modalities [9]. Crucially, canine oral MM occurs at higher frequency and progresses over a shorter timeframe than human MM: canine patients survive for a median of just 335 days following surgical excision and systemic adjuvant therapy [9,10].

Previous research has identified two transcriptomic subgroups in primary OMM in both dogs and humans [8]. These subtypes reveal, for the first time, heterogeneity in the immune microenvironment and point toward potential therapeutic targets, but they have not been definitively linked to prognosis. In both species, regional lymph node metastasis is common, yet no prior study has examined the molecular characteristics of these nodes to assess their prognostic value or therapeutic implications. A deeper understanding of the genetic and transcriptomic landscapes of both metastatic and non-metastatic lymph nodes could offer valuable prognostic insights at the time of diagnosis and inform precision medicine approaches [3,11–16].

We have curated a cohort of formalin-fixed, paraffin embedded samples of naturally-occurring, treatment-naïve, OMM from canine patients [8]. This collection is complemented by comprehensive clinicopathological information sourced from clinical records. Here, for the first time, we apply transcriptomic data from surgically excised regional lymph nodes of dogs with OMM. We demonstrate that lymph nodes can be stratified into two distinct transcriptomic subgroups, independent of their histopathological metastatic status. This stratification can be succinctly captured using a 35-gene signature. These subgroups reflect immune “hot” and “cold” states, correlate with patient overall survival, and provide clinically meaningful stratification at the time of OMM diagnosis. Finally, we identify IBA1 immunohistochemistry as a practical biomarker to stratify lymph node states, offering a clinically actionable tool to refine prognosis and inform the design of future clinical trials.

## MATERIALS AND METHODS

### EXPERIMENTAL MODEL AND STUDY PARTICIPANT DETAILS

#### Ethics Statements

This study was conducted with the approval of the University of Edinburgh’s Veterinary Ethical Review Committee (Reference number 10.21). Tissue was only included in the study with the informed, written consent of the caregiver of the dog who bore the tumour. The treatment that a canine patient received was unaffected by their inclusion in this study.

De-identified canine clinicopathological, genetic and standardised RNA sequencing data have been deposited at the Sequence Read Archive (SRA with accession number #SUB14547907).

##### Sample collection

Formalin-fixed, paraffin-embedded (FFPE) canine OMM tissue samples and associated clinical data were collected from patients with naturally occurring tumours who underwent surgical resection or biopsy of the primary tumour (as previously described by Bowlt Blacklock et al [8]).

FFPE tissue samples of treatment-naïve canine OMMs were collected between 2011 and 2021 for histopathology from dogs attending the Clinical Veterinary Oncology departments at the Universities of Edinburgh, Bristol, Liverpool, Glasgow and Ohio Small Animal Teaching Hospitals, Vets Now (Glasgow) Referrals, Paragon Veterinary Referrals, The Ralph Veterinary Referral Centre, Anderson Moores Veterinary Specialists, and North Downs Specialist Referrals. Clinicopathological data collected included sex, breed, age (years), primary tumour anatomical location, metastatic status (confirmed by abdominal ultrasound, thoracic computed tomography or radiography, and/or histological examination of ≥1 regional lymph node), World Health Organization (WHO) status, provision of adjuvant therapy, and duration of patient survival or date lost to follow up (in months following initial diagnosis of OMM).

The diagnosis of primary OMM was confirmed by a board-certified veterinary anatomic pathologist (JdP). Lymph nodes were classified as histopathologically metastatic (MT) or non-metastatic (NMT). Reflecting real-world diagnostic practice, the histopathological metastatic status of the regional lymph nodes was assessed using haematoxylin and eosin (H&E) staining. In cases where morphology was equivocal, routine immunohistochemistry (IHC; Melan-A, PLN2, and/or Iba1) was performed at the discretion of the pathologist and as part of the clinical work-up to support diagnosis. Lymph nodes with sparse or early metastatic features were classified as metastatic. Where metastasis was histopathologically apparent, representative tumour regions were identified on FFPE blocks.

Accurate attribution of cause of death in this population was challenging. Unlike in human oncology, death certification is not routinely available for veterinary patients, and most dogs were euthanised rather than dying naturally. In addition, clinical follow-up was split between referral hospitals and primary care practices, with variable detail available in the medical record. Only four patients had a documented cause of death, all of whom were seen at referral hospitals for an unrelated condition at the time of euthanasia. For the remainder, it was generally not possible to distinguish whether death was directly attributable to melanoma progression or to other causes.

## METHOD DETAILS

### RNA extraction

Using a manual tissue arrayer (Beecher MTA-1, Estigen OÜ, Estonia) machine, two cores (1mm outer diameter, 5mm length) were harvested from each canine FFPE block: one from a representative area of lymph node tissue and one from histopathologically normal tissue. For nodes classified as metastatic, the core was taken from a region containing tumour cells. For nodes classified as non-metastatic, the core was procured from a representative area of nodal tissue. Normal tissue core was obtained from an unaffected area within the FFPE block containing the primary tumour. This approach ensured all samples included both a representative lymph node core and a comparator core from non-tumour tissue. Tissue selection was guided by a board-certified anatomic pathologist (JdP) to minimise tissue heterogeneity and avoid sampling bias. RNA was isolated from the tissue cores using the Covaris E220 Evolution Focused Ultrasonicator (Covaris Inc, #500429) and the truXTRAC® FFPE total NA (Nucleic Acid) Kit – Column (Covaris, #520220). Inhibitors (e.g. melanin) were removed using the OneStep PCR Inhibitor Removal kit (Zymo Research #D6030).

#### Quality control

Total RNA was assessed for quality and integrity on the Fragment Analyser Automated Capillary Electrophoresis System (Agilent Technologies Inc, #5300) with the Standard Sensitivity RNA Analysis Kit (#DNF-471-0500), and then quantified using the Qubit 2.0 Fluorometer (Thermo Fisher Scientific Inc, #Q32866) and the Qubit RNA (#Q10210) Broad Range assay kits. DNA contamination was quantified using the Qubit dsDNA HS assay kit (#Q32854). The Fragment Analyser data showed a high degree of RNA degradation so the fragmentation step was omitted from the library preparation protocol.

#### RNA library preparation

First-strand cDNA was generated from 50 ng of each total RNA sample using the SMARTer® Stranded Total RNA-Seq Kit v2 – Pico Input Mammalian kit (Clontech Laboratories, Inc. #634411). Illumina-compatible adapters and indexes were then added via 5 cycles of PCR. AMPure XP beads (Beckman Coulter, #A63881) were then used to purify the cDNA library. Depletion of ribosomal cDNA (cDNA fragments originating from highly abundant rRNA molecules) was performed using ZapR v2 and R-probes v2 specific to mammalian ribosomal RNA. Uncleaved fragments were then enriched by 13 cycles of PCR before a final library purification using AMPure XP beads. Libraries were quantified with the Qubit dsDNA HS assay and assessed for quality and size distribution of library fragments using the Fragment Analyser and the NGS Fragment Kit (#DNF-473-0500).

#### Sequencing

Sequencing was performed on the NextSeq 2000 platform (Illumina Inc, #20038897) using NextSeq 2000 P3 Reagents (200 Cycles) (#20040560). Libraries were combined in equimolar pools based on Qubit and Bioanalyser assay results and each pool was sequenced on a P3 flow cell. For RNA sequencing, PhiX Control v3 (Illumina Inc, #FC-110-3001) was spiked into each run at a concentration of 1% to allow troubleshooting in the event of any issues. Basecall data produced by the NextSeq 1000/2000 Control Software (Version 1.4.1.39716) was automatically converted into FASTQ files and uploaded to BaseSpace.

### Immunohistochemistry

Immunohistochemistry was conducted on 4µm-thick formalin-fixed paraffin-embedded canine OMM tissue, acquired using a microtome (Thermo Scientific Rotary Microtome Microm HM 340 E). All tissue sections, including relevant controls, were deparaffinized in xylene and rehydrated in graded ethanol through to distilled H_2_0 prior to staining. Heat-induced antigen retrieval was performed using sodium citrate (pH 6.0; 110°C; 5-12 minutes overall) in a microwave (HistoS5, Milestone). Once retrieved, all tissue sections were stained in relevant batches in an autostainer (Epredia Autostainer 360) following a standard protocol which incorporates a 30-minute incubation with the primary antibody at room temperature. CD3 was used to identify T cells and NK cells (Novocastra Liquid Mouse Monoclonal Antibody CD3, Leica Biosystems, #NCL-L-CD3-565; 1/200 dilution), PAX5 was employed to stain B lymphocytes (Mouse Monoclonal Antibody Pax5, Becton and Dickinson, P67320-050; 1/50 dilution), and anti-IBA1 was used to label monocytes/macrophages (Rabbit polyclonal anti Iba1, Wako, 019-19741; 1/500 dilution). All sections were treated with Dako Real Peroxidase Blocking solution (Dako S2023) to block against endogenous peroxidase. The next step was a 40-minute incubation with either EnVision anti-mouse (Dako, EnVision+ System –HRP Labelled Polymer; 302059EFG_00) or EnVision Anti-Rabbit. (Dako, EnVision+ System HRP-labelled polymer; K400311-2). This system is based on an HRP labelled polymer which is conjugated with secondary antibodies. The labelled polymer does not contain avidin or biotin. As such, nonspecific staining resulting from endogenous avidin-biotin activity is eliminated or significantly reduced. EnVision mouse was used to stain tissues with CD3 and PAX5, whereas EnVision Anti-Rabbit with DAB enhancer protocol was used for IBA1 (Dako EnVision®+ Dual Link System-HRP (DAB+), PD04048_02/K4065). All sections were then counterstained with Harris haematoxylin (Varistain Gemini Slide Stainer, Thermo Fischer Scientific), dehydrated with graded ethanol through to xylene, and covered with coverslips.

### Image acquisition

Slides were digitally scanned (Nanozoomer-XR, Hamamtsu Photonics K.K, Japan) and raw image data were saved in .ndpi format and handled by the software ndp.view 2 (Hamamatsu Photonics) to save details of the whole image in .jpeg format. Images were imported to ImageJ (FiJi Project, 2.11.0, GPLv3+) for labelling, and uploaded to QuPath (v0.5.0)[17].

## QUANTIFICATION AND STATISTICAL ANALYSIS

### Data analysis pipeline

#### Alignment and Gene-Level Counts

RNA-seq data were processed using the nf-core ’rnaseq’ pipeline v3.8.1[18–32]. In brief, samples were aligned via STAR v2.6.1d, and gene-based counts produced using Salmon v.1.5.2. Canine data were aligned to the ROS_Cfam_1.0 reference genome, annotated using the corresponding GTF file for build accession GCA_014441545.1.

##### Unsupervised Clustering

Unsupervised consensus clustering of expression data was performed using the R Bioconductor package, ’cola’ [33]. Prior to clustering, gene level count data were subject to a variance stabilising transformation using DESeq2 [29], and the resulting matrix was used as an input. Five different clustering algorithms were tested, evaluating two to six clusters in each case. The cola algorithm resamples count data a fixed number of times, repeating the clustering process on each iteration. The optimum clustering strategy was selected on the stability of the resulting clusters; the method by which samples cluster most consistently.

##### Differential Expression Analysis

Differential expression analysis was performed using DESeq2. Models were fitted treating the clusters identified by cola, sex, and, where appropriate, batch as factors. Log fold change (logFC) estimates were produced using the apeglm shrinkage method [34], which is intended provide more robust estimates in the event of high within-group variability. Shrunken logFC estimates were accompanied by s-values [35], an aggregate false sign rate which are broadly analogous to q-values. P-values were also computed and adjusted using the IHW method [36]. Results were annotated using biomaRt [37], and volcano plots generated using EnhancedVolcano [38].

##### Over-Representation Analysis

Over-representation analysis (ORA) was performed via the R package ClusterProfiler[39]. ORA was performed for up- and down-regulated genes independently, taking the top differentially expressed genes with an s-value < 0.01 in each case. These gene lists were then tested for over-representation against the Biological Process (BP), Cellular Component (CC), and Molecular Function GO ontologies [40], and in addition to known pathways from the Kyoto Encyclopedia of Genes and Genomes (KEGG) [41]. Minimum and maximum gene set sizes were set to 10 and 1,000, respectively for all analyses.

#### Immunocyte deconvolution

The computational framework CIBERSORTx was used to deconvolute the tumour-infiltrating immune cells from bulk RNA-sequencing data [42]. Twenty-two immune cell subtypes were parsed from the annotated gene signature matrix LM22 and 100 permutations of the CIBERSORTx web portal [42]. After running, only samples with CIBERSORT *p*-values < 0.05 were included in subsequent analyses. B-mode batch correction was applied, and quantile normalisation was disabled to generate absolute scores. CIBERSORT fractions for naïve B cells, memory B cells and plasma cells were aggregated into B cells; monocytes, M0, M1, M2 macrophages into monocytes/macrophages; and CD8, CD4, T follicular helper cells, regulatory T cells, and gamma delta T cells into T cells; and resting and activated NK cells into NK cells. Wilcoxon tests were used to compare the immune cell fractions between transcriptomic consensus clustering groups. A cox-regression model was used to describe the time-to-event outcomes, with the adjustedCurves package used [43] to show the effect of a continuous variable adjusted for age and gender/sex. Receiver Operator Curves were plotted and optimal cut off points were defined using Youden’s index. Statistical significance was set at p<0.05.

#### Image data analysis

Using the H&E slides to identify the tumour margins, the tumour areas were defined in QuPath. Positive cell detection was performed using the default parameters, except for sigma, which was set at 1.0μm. Wilcoxon tests were used to compare the ‘number of positive cells detected per mm^2^’ between transcriptomic consensus clustering groups. Optimal cut off points were defined as above. Statistical significance was set at p<0.05.

#### Results

We identified canine (Bowlt Blacklock cohort [8]; n=37) patients who presented for treatment of naturally occurring, treatment-naïve OMM and had archival formalin-fixed-paraffin-embedded (FFPE) tissue and clinical records available for analysis (**Figure 1a)**. Dog and tumour characteristics are summarised in **Supplementary Data 1**, as previously described by Bowlt Blacklock *et al* [8].

**Figure 1:**
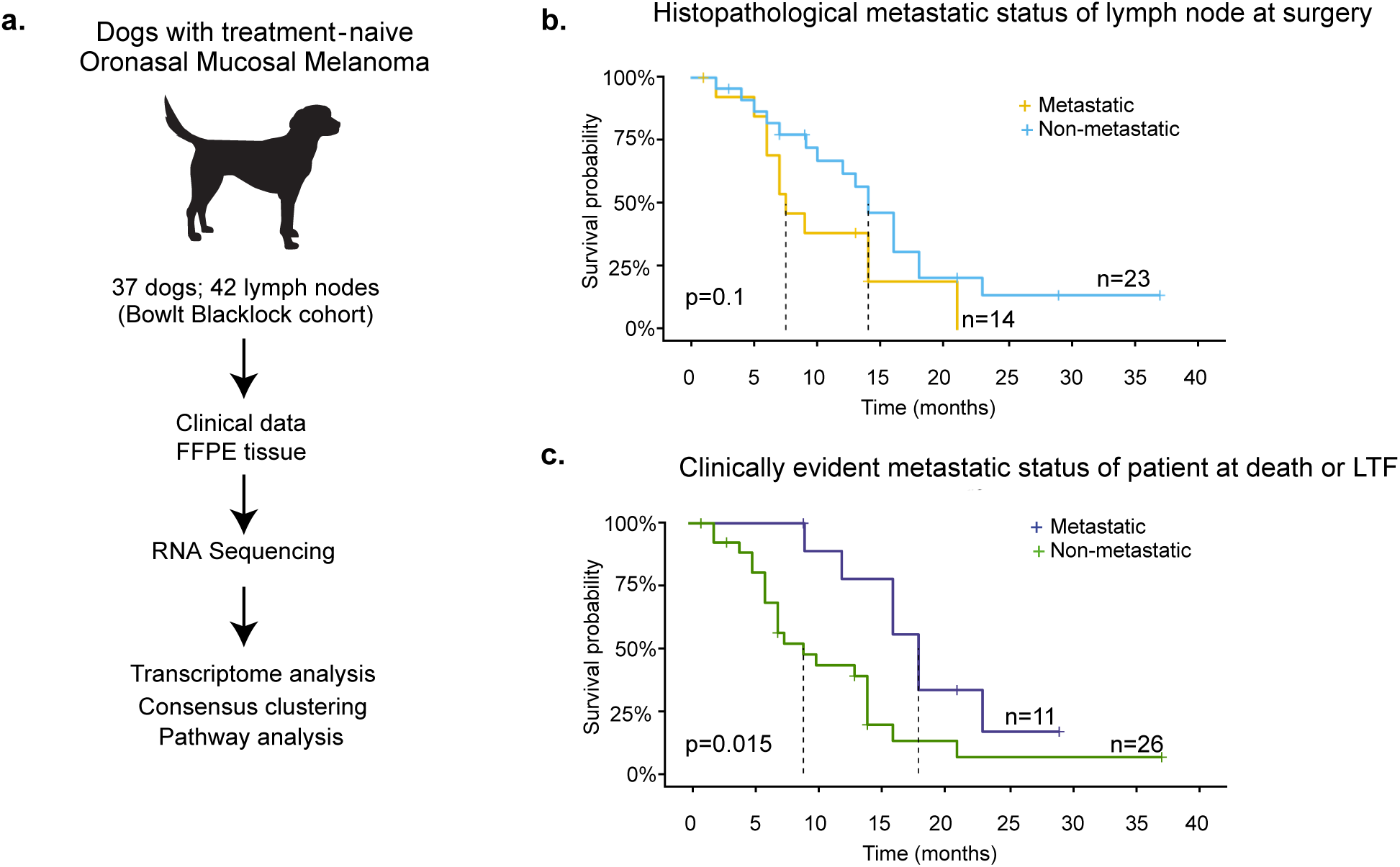
Clinical metastatic status at the time of diagnosis does not predict survival. **(A)** Project workflow. Dogs with treatment-naïve oronasal mucosal melanoma (n = 37) underwent surgical resection of the primary tumour (n=37) and regional lymph nodes (n=42). FFPE tissues were used for transcriptomic analyses. **(B)** Kaplan-Meier survival plot comparing dogs with and without histologically confirmed lymph node metastasis at the time of OMM diagnosis. **(C)** Kaplan-Meier survival plot comparing dogs with and without clinically evident metastatic disease (lymph node and/or distant) recorded at death or loss to follow-up (LTF). Clinical evidence of metastasis was determined by staging investigations (thoracic/abdominal/head imaging) or by detection of progressive nodal metastases on clinical examination. Dotted horizontal lines represent the 50% survival probability threshold in each plot.

Forty-two lymph nodes were surgically extirpated at the same time as resection of the primary tumour: 34 patients had 1 node removed, 2 patients had 2 nodes removed, and 1 patient had 4 nodes removed; 14 nodes from 14 dogs were histopathologically consistent with metastatic disease, including both overt and early metastasis as determined by a board-certified pathologist. At the time of death or loss to follow-up, 24 dogs had developed nodal or distant metastatic disease.

To avoid biological pseudo-replication, only one lymph node sample per dog was included in the differential expression and clustering analyses. In cases where both metastatic (MT) and non-metastatic (NMT) nodes were available from a single patient, the metastatic node was prioritised for inclusion. This *a priori* decision reduced intra-individual bias, ensured balanced representation across groups, and avoided inflating sample size. In total, only five NMT samples from three dogs were excluded from analysis.

##### Lymph node metastatic status at the time of diagnosis does not predict survival

Given that lymph node involvement is commonly used as an indicator of disease progression, we assessed whether the histopathological metastatic status of surgically removed nodes was predictive of survival. No statistically significant difference in overall survival was observed between dogs with or without histopathological lymph node metastasis at the time of OMM diagnosis (p = 0.1; **Figure 1b).** However, patients who developed clinically evident metastatic disease at any time during follow-up had significantly worse survival outcomes (p = 0.015; **Figure 1c).** Across the whole cohort, long-term survival remained poor overall.

Our data indicate that lymph node metastatic status, as assessed histopathologically at the time of surgery, does not reliably predict survival and therefore has limited prognostic value at the time of diagnosis. Given this limitation, we next explored whether transcriptomic profiling of both metastatic and non-metastatic lymph nodes could reveal molecular features with greater prognostic or biological relevance.

##### Lymph node transcriptomic subgroups outperform histopathology in prognostication

When transcriptomic data from all lymph nodes—regardless of histopathological metastatic status—were analysed, we identified two stable consensus clusters (A and B) using the optimal consensus partitioning method with standard deviation and spherical k-means clustering (**Supplementary Data 2**). Principal component analysis showed distinct grouping of the two clusters along PC1 (**Figure 2a**).

**Figure 2:**
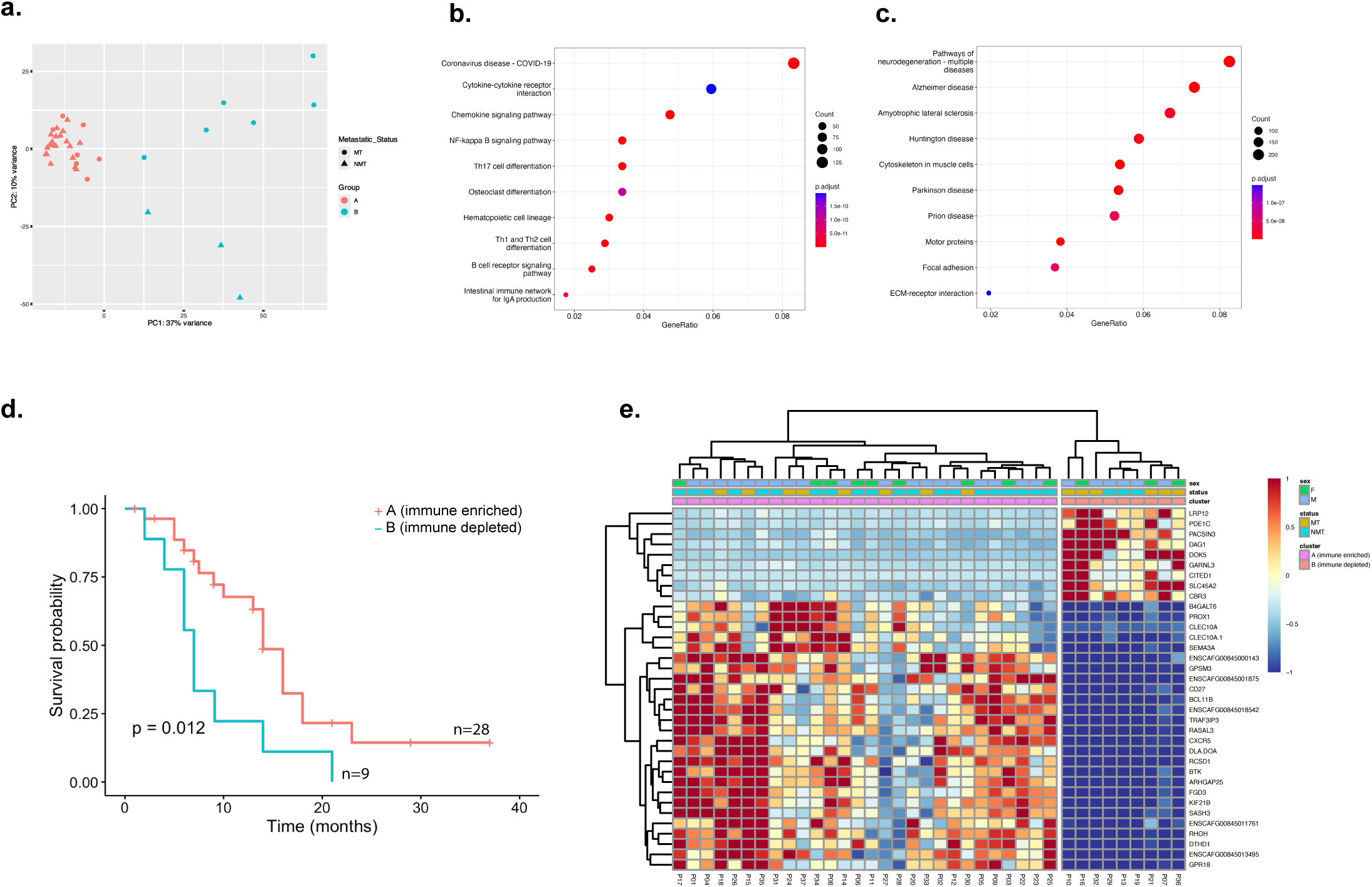
Lymph node transcriptomic signatures predict survival in canine OMM. **(A)** Principal component analysis of all lymph node transcriptomes identified two distinct and stable clusters, independent of histopathological metastatic status at the time of surgery: subgroup A (“immune-enriched”) and subgroup B (“immune-depleted”). **(B–C)** Over-representation analysis (ORA) of KEGG pathways revealed that the ‘immune-enriched’ subgroup (B) was enriched for immune-related pathways, while the ‘immune-depleted’ subgroup (C) showed enrichment for neurodegenerative signalling pathways. **(D)** Kaplan–Meier survival curve comparing dogs by transcriptomic subgroup. **(E)** Heatmap showing expression of a 35-gene classifier identified using random forest modelling that stratified lymph node samples into transcriptomic subgroups.

To understand the pathways that characterised the transcriptional subtypes, we applied over-representation analysis (ORA) and showed that pathways align with immune pathways in subgroup A (**Figure 2b**) and neurodegenerative responses in subgroup B (**Figure 2c**).

We examined whether the transcriptomic subgroups were associated with clinical or pathological features, including age, sex, tumour site, breed, and primary tumour pigmentation; however, none of these comparisons reached statistical significance.

We then asked whether the transcriptomic subgroups simply mirrored histopathological metastatic status at diagnosis. Principal component analysis showed partial separation between MT and NMT nodes, but with substantial overlap suggestive of shared molecular traits or intermediate states (**Figure 2a**). This overlap may reflect occult metastatic disease not diagnosed by histopathology, but could equally indicate nodes actively mounting an immune response that prevents disease progression. Importantly, lymph node assignment to a transcriptomic subgroup was independent of both histopathological status at the time of surgery (Fisher’s exact test, OR=0.20, p=0.06), and clinically evident metastatic status at patient death or loss-to-follow up (Fisher’s exact test, OR=3.79, p=0.39), (**Figure 2a; Supplementary Data 2**). This held true even for dogs whose nodes were initially deemed non-metastatic but who later developed clinical evidence of metastases (13 of 23 dogs), as such patients were distributed across both transcriptomic subgroups (**Supplementary Data 2**).

We next examined whether transcriptomic subgroup classification was associated with patient survival. Patients with lymph nodes assigned to subgroup A had a statistically improved overall survival (median 13 months, range 1–37 months) compared with those in subgroup B (median 7 months, range 2–21 months) (p=0.012) (**Figure 2d**). All patients lost-to-follow-up (n=6) were in subgroup A, and time to loss to follow up was similar to known survival time (median time to loss to follow up 11 months, mean 15.8 months, range 1-37 months).

As an internal check, we analysed MT and NMT nodes separately. While each group formed distinct transcriptomic clusters, these within-group clusters were not prognostic (**Supplementary Figure S1**). This confirmed that prognostic information is most effectively captured when all lymph nodes are analysed together, irrespective of histological classification at the time of diagnosis, and that transcriptomic subgroups capture biology beyond MT/NMT classification.

Finally, we aimed to find the most important variables that drive the stratification across cohorts. First, we filtered the dataset for the top 3000 most variably expressed genes, which are consistently detected in at least 10% of samples. Next, we used the machine learning algorithm random Forest [44] across three randomly selected seeds to generate successful classification models (trained on 50% of the samples) that predicted subgroup B versus the A subgroup in the test data (the remaining 50% of the samples), independent of metastatic status or sex information (accuracy >90%, binomial test, p-value <0.05). Within these models, 35 genes were consistently ranked among the most important genes shared by all three models (relative variable importance value >1%) (**Supplementary Data 3, Figure 2e)**.

Using g:Profiler [45,46], we mapped the 35-gene lists to functional information sources to detect statistically significantly enriched terms (**Figure 3e)**. Lymph nodes assigned to the subgroup B were aligned with increased expression of genes associated with melanin biosynthesis. By contrast, lymph nodes assigned to the subgroup A were associated with pathways related to lymphocyte activation (**Supplementary Data 3)**.

**Figure 3:**
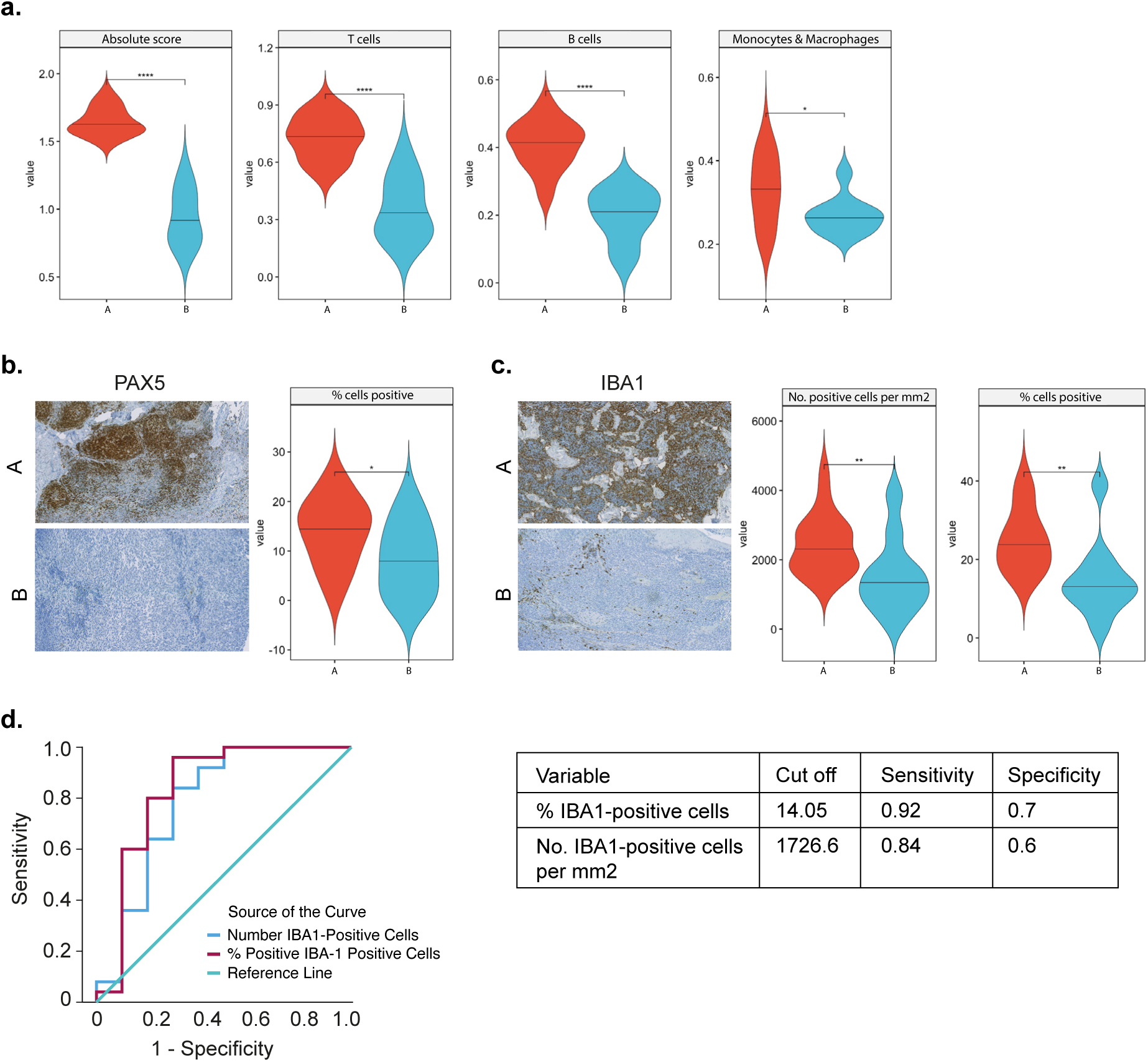
Monocytes/macrophages and B-cells are diagnostic markers for transcriptomic subgroups in canine OMM lymph nodes. **(A)** Violin plots for immunocyte infiltration of canine OMM lymph nodes, determined by CIBERSORTx, stratified according to transcriptomic subgroup*. **(B)** Representative IHC panel and violin plot of PAX5+ B-cell infiltration, stratified according to transcriptomic subgroup*. **(C)** Representative IHC panel and violin plot of IBA1+ monocyte/macrophage infiltration, stratified according to transcriptomic subgroup*. **(D)** Receiver operating characteristic (ROC) curves and optimal cut-off thresholds for B-cell and monocyte/macrophage infiltration as diagnostic classifiers of lymph node transcriptomic subgroup in canine OMM. * subgroup A: “immune-enriched”; subgroup B :”immune-depleted”.

Collectively, these findings indicate that transcriptomic subgroups provide prognostic insights beyond conventional metastatic status, suggesting that draining lymph nodes act as active integrators of tumour–immune signals. Our data suggests that lymph nodes that mounts an immune response (subgroup A) support improved clinical outcomes for patients, while dogs with lymph nodes with a relatively poor immune response at time of surgery (subgroup B) have poorer ourcomes.

##### Tumour-educated lymph nodes offer improved prognostication when compared to the primary tumour

Primary OMM tumour transcriptomes from the same dogs included in this study have previously been classified into two subgroups: CTLA4-high (immune ‘hot’) or MET-high (immune ‘cold’) [47]. However, unlike the lymph-node–based subgroups, these primary-tumour-derived classifications did not correlate with survival outcomes. Moreover, there was no correlation between primary tumour classification and lymph node subgroup assignment (Fisher’s exact test, OR=1.6, p=0.71)**(Supplementary data 2)**.

This represents a key conceptual advance: lymph nodes may act as early biosensors of tumour-derived signals, independent of their histopathological status.

##### IBA1 Demonstrates Strong Diagnostic Potential for Lymph Node Stratification

The identification of two transcriptomic subgroups suggested that patient stratification could inform prognosis and guide therapeutic decisions or clinical trial design. Because transcriptomic profiling is not currently practical in most clinical settings, we therefore sought diagnostic immunohistochemical (IHC) markers to distinguish subtypes.

We hypothesised that subgroup A could be identified by immune infiltrates and used CIBERSORTx [42] to infer cell composition from bulk RNA-seq data (**Supplementary Data 4**). Across all lymph nodes, subgroup A showed significantly greater total immune infiltration (Absolute Score, p<0.01), including higher levels of T cells, B cells, monocytes, and macrophages (**Figure 3a**).Therefore, we named subgroup A as ‘immune-enriched’, and subgroup B as ‘immune-depleted’.

To test whether these transcriptomic signals could translate into a diagnostic assay, we applied IHC with digital pathology [17] to characterise immune infiltrates across whole lymph nodes using markers for T cells (CD3), B cells (PAX5), and monocytes/macrophages (IBA1). CD3+ T cells were more abundant in the ‘immune-enriched’ subgroup but not significantly (p = 0.101). PAX5+ B cells were significantly higher in ‘immune-enriched’ nodes (p=0.037; **Figure 3b**). The largest differences were observed for IBA1+ monocytes/macrophages, with both the percentage of positive cells (p=0.001) and number of positive cells per mm² (p=0.008) significantly elevated in ‘immune-enriched’ nodes (**Figure 3c**).

Receiver Operator Curve (ROC) analysis was used to evaluate the diagnostic performance of PAX5 and IBA1. Using the Youden index, we defined clinical relevance as an AUC (area under the curve) >0.75. PAX5 (% positive cells, AUC=0.73) did not meet this threshold. By contrast, IBA1 variables showed strong discriminatory power, with an AUC of 0.84 for % positive cells and 0.78 for number of positive cells per mm^2^ (**Figure 3d**). A threshold of 14.05% IBA1-positive cells discriminated between subgroups with high diagnostic performance (sensitivity 0.92, specificity 0.70).

These results suggest that ‘‘immune-enriched’ lymph nodes (subgroup A) represent an “immune-hot” microenvironment, characterised by abundant macrophage infiltration, whereas ‘immune-depleted’ nodes (subgroup B) reflect a comparatively “immune-cold” state. This distinction highlights IBA1 as a clinically relevant marker for stratifying lymph node subgroups, providing a meaningful tool to refine prognostic evaluation in canine OMM.

##### DNA tyrosinase vaccine does not improve survival across stratified subgroups

Finally, to assess the potential clinical impact of existing immunotherapies in stratified canine patients, we analysed outcomes following administration of a xenogeneic (human) DNA vaccine targeting tyrosinase (Oncept). No significant difference in overall survival was observed between vaccinated and unvaccinated dogs, regardless of nodal histopathological status (metastatic or non-metastatic) or transcriptomic subgroup (‘immune-enriched’ or ‘immune-depleted’) (**Supplementary Figure S2**).

Thus, while Oncept has been used in stage II–III canine oral melanoma and reportedly offers some survival benefit in uncontrolled studies [48], our stratified analysis still showed no survival difference. These findings underscore the need to expand immunotherapeutic strategies in OMM and reinforce the value of biomarker-guided approaches to enable more effective, targeted interventions.

## Discussion

In this study, we provide the first transcriptomic analysis of regional lymph nodes in dogs with OMM. We demonstrate that nodes can be stratified into two distinct subgroups, independent of histopathological metastatic status, and that this stratification correlates with patient survival. Importantly, these transcriptomic profiles capture prognostic information not evident from conventional histopathology, highlighting their potential to refine risk assessment at the time of diagnosis.

Our findings underscore a fundamental limitation of current staging practices: histopathological assessment of lymph nodes, while widely used at the time of surgery, does not reliably predict outcome. This may in part reflect the inherent limitations of morphological assessment, where micrometastatic disease can be missed due to sampling bias or the restricted sensitivity of routine H&E and IHC. Transcriptomic profiling may therefore reveal occult metastatic or pre-metastatic niches that are not morphologically detectable, providing a more comprehensive readout of tumour–immune dynamics [49–57].

The biological basis of the two subgroups is informative. ‘Immune-enriched’ nodes, associated with longer survival, were characterised by “immune-hot” features, including increased infiltration of monocytes/macrophages and B cells, as well as enrichment of immune-related pathways. By contrast, ‘immune-depleted’ nodes displayed an “immune-cold” phenotype underpinned by neurodegenerative and stress-response pathways. This suggests that lymph nodes act as biosensors, integrating tumour-derived signals and host immune responses in ways that transcend their histological classification. It has been demonstrated that sentinel lymph nodes from human melanoma patients are stratified along a naïve to activated immune axis, and dendritic cell infiltration is also a key component of antitumour immune response [58]. Notably, these immune-based phenotypes were not recapitulated by transcriptomic subgroups of the primary tumour, reinforcing the concept that nodes may provide sensitive prognostic information that is not currently detectable in the primary tumour itself. In human melanoma, a “hot” immune microenvironment has been correlated with better outcomes and greater sensitivity to immune-based interventions [59]. Similarly, in canine apocrine gland anal sac adenocarcinoma (AGASACA), immune “hot” tumours correlate with smaller tumour size and absence of metastasis [60]. In contrast, tumours with an immune-excluded or immune-suppressed phenotype often exhibit resistance to immunotherapy due to mechanisms such as T cell exhaustion, myeloid-derived suppressor cell (MDSC) expansion, or an immunosuppressive cytokine milieu [61–63]. Instinctively, our ‘immune-enriched’ subgroup may benefit from immunotherapeutic strategies, and identifying and characterising immune-enriched lymph node subgroups could serve as a crucial step in stratifying patients for immunotherapy, an emerging treatment option in dogs [63,66].

In contrast, the ‘immune-depleted’ subgroup may require alternative approaches to overcome immune evasion. Tumours exhibiting an immune-excluded phenotype often evade immune surveillance by suppressing cytotoxic lymphocyte infiltration, upregulating inhibitory signals, or fostering a tumour-promoting stromal environment [67]. In such cases, conventional immunotherapies may be ineffective due to a lack of pre-existing immune activation.

Although xenogeneic (human) DNA vaccination targeting tyrosinase (Oncept) is licensed for canine oral melanoma [68], we found no survival benefit in dogs receiving the vaccine across any stratified subgroup. This may be because the vaccine may be administered too late to prevent micrometastatic progression, particularly in dogs with established or biologically aggressive disease [69,70]. These findings underscore the limitations of current non-stratified immunotherapeutic approaches in OMM and highlight the need for biomarker-guided strategies. While xenogeneic vaccines such as tyrosinase-targeted Oncept have not shown clear benefit in our cohort, antigen-based therapies remain a promising avenue. The key challenge will be in identifying the most relevant antigens and aligning them with the immune contexture of individual patients to optimise efficacy.

From a translational perspective, we identified IBA1+ monocyte/macrophage infiltration as a robust biomarker of transcriptomic subgroups. While RNA sequencing is unlikely to be widely feasible in routine clinical workflows, IBA1 immunohistochemistry is scalable, reproducible, and can be readily incorporated into standard pathology practice [71]. Thus, IBA1 has strong potential as a clinically actionable biomarker to stratify patients, refine prognosis, and guide therapeutic decision-making.

Our study has several limitations. The sample size, while robust for a naturally occurring veterinary cohort, is small and should be expanded to confirm the generalisability of our results. Additionally, while transcriptomic profiling provides valuable biological insights, further functional validation of the identified pathways is warranted. The lymph nodes analysed in this study were identified for removal on clinical grounds, and were not confirmed as sentinel nodes, which may have resulted in underdiagnosis of metastatic disease in patients where the examined node was not the primary lymphatic drainage site [72]. It should be noted that clinically evident metastatic disease status during patient follow-up was determined based on clinical need rather than systematic longitudinal assessment. Follow-up investigations varied (e.g. imaging, fine needle aspiration, clinical notes), and therefore subclinical metastatic disease may have been missed. In addition, some patients may have died of unrelated causes, before metastatic progression could be documented. Unlike in human oncology, death certification is not routinely available for veterinary patients, and most dogs were euthanased rather than dying.

Standard histopathological examination differentiated metastatic from non-metastatic lymph nodes, but is neither 100% sensitive or specific, as early metastases may be missed or melanocytic cells misclassified as metastatic cells [73,74]. Immunohistochemistry or in situ hybridisation would enhance accuracy in differentiating true metastases, but is uncommonly performed in a clinical setting [74–77]. While tumour burden within lymph node cores may vary, the robust correlation of transcriptomic clusters with survival outcomes and IBA1 expression supports the biological relevance of the identified subgroups beyond sampling differences.

Exploring whether the identified subgroups in canine melanoma extend to human mucosal melanoma holds significant promise for cross-species translational research. If specific molecular or immunological subtypes are conserved between dogs and humans, then therapies showing efficacy in canine models might be more readily translated into human clinical trials [78]. Longitudinal transcriptomic data collection from primary tumours and metastatic nodes may identify prognostic biomarkers, and guide adaptive therapies for better survival in both species.

Transcriptomic analysis can be carried out on fine needle aspirate samples from some tumours [79–81] and a major clinical consideration is whether the identified transcriptomic signatures can be reliably detected in fine needle aspirates of lymph nodes [82]. If these molecular signatures can effectively stratify patients, it may prompt review of decision making in both human and canine oncology. In human oncology, the benefits of prophylactic lymph node dissection are increasingly questioned [83], and similar considerations may shape the management of canine OMM [84]

### Conceptual Advance and Translational Potential

The conceptual advance in our work is the discovery that lymph nodes from canine OMM can be stratified into two transcriptomic subgroups that correlate with patient survival. Patients with ‘immune-enriched’ lymph nodes, with increased infiltration of B cells, monocytes, and macrophages, have improved outcomes compared with patients with ‘immune-depleted’ lymph nodes, underpinned by neurodegenerative and stress-response pathways. These findings suggest that patients with ‘immune-enriched’ lymph nodes may be more likely to benefit from immunotherapy, whereas those with ‘immune-depleted’ nodes may respond better to targeted strategies. Biologically, this highlights the draining lymph node as a gatekeeper of disease progression: when an effective immune response is mounted, metastatic spread may be delayed and survival prolonged, whereas failure to generate such a response permits dissemination and poor outcome. Because human and canine OMM are fundamentally the same disease, these insights extend beyond companion animal oncology and offer a translational model for understanding immune containment and progression in human mucosal melanoma.

## Acknowledgements

We extend our heartfelt gratitude to all the patients or caregivers who generously consented to contribute samples for this study. The authors thank the following colleagues for their expertise and assistance in this work: Angie Fawkes, Lee Murphy and Richard Clark (Edinburgh Clinical Research Facility, University of Edinburgh), Steve Brawley (Vets4Pets), and Helen Caldwell (University of Edinburgh). This work was funded by the Kennel Club Charitable Trust (10915550_10917251), Melanoma Research Alliance and Rosetrees trust (687306), Cancer Research UK Scotland (CTRQQR-2021), and the Medical Research Council (MC_UU_0035/13).

## Author Contributions Statement

KLBB and EEP conceptualised and designed the study. Data collection was performed by AL, LB, KLBB, JdP, GP, LS, J-BT, DK, JSM, IB, SZ, SMG, MST, DM, AM, KP and MC. Data analysis was conducted by KD, YL, AL, AM, EEP and KLBB. Data interpretation was contributed by LB, AL, KD, YL, KLBB and EEP. KLBB, LB, AL and EEP conducted the literature search and generated the figures. All authors were involved in writing the paper and had final approval of the submitted and published versions. KLBB provided supervision and project administration.

## Data availability statement

De-identified canine clinicopathological and standardised RNA-seq data were deposited at the Sequence Read Archive (SRA with accession no. SUB14547907).

**Supplementary Figure S1. Transcriptomic variation within histopathologically classified lymph nodes reveals biological divergence without survival association**

**(A)** Principal component analysis (PCA) of transcriptomic data from metastatic (MT) nodes only (left), and non-metastatic (NMT) nodes only (right). Distinct subclusters (MT-A/MT-B and NMT-A/NMT-B) were identified using unsupervised consensus clustering.

**(B)** Over-representation analysis (ORA) of KEGG pathways in MT nodes revealed enrichment for immune-associated pathways in MT-A (left), and neurodegenerative or structural signalling pathways in MT-B (right), suggesting divergent biological programs despite shared histopathological classification. Optimal clustering was performed using mean absolute deviation and spherical k-means methods.

**(C)** Over-representation analysis (ORA) of KEGG pathways in NMT nodes revealed enrichment for immune-associated pathways in NMT-A (left), and cellular or structural signalling pathways in MT-B (right), suggesting microenvironmental activation or tissue response processes in histologically non-metastatic nodes. Optimal clustering was performed using standard deviation and spherical k-means methods.

**(D)** Kaplan–Meier survival curves comparing MT-A and MT-B subgroups (left) and NMT-A and NMT-B subgroups (right). No significant differences in survival were observed, suggesting that the observed transcriptomic divergence does not correlate with clinical outcome in this cohort.

**Supplementary Figure S2. Survival outcomes following human DNA tyrosinase (Oncept) vaccination do not differ by nodal metastatic status or transcriptomic subgroup**

(A) Kaplan–Meier survival plot comparing vaccinated and unvaccinated dogs with histologically confirmed metastatic and non-metastatic lymph nodes. No significant difference in survival was observed between groups.

(B) Kaplan–Meier survival plot comparing vaccinated and unvaccinated dogs assigned to transcriptomic subgroup A (“immune-enriched”) or B (“immune-depleted”). Oncept vaccination did not significantly alter survival outcomes in either group.

(C) Kaplan-Meier survival plot of vaccinated dogs with OMM, stratified by transcriptomic subgroup (A ‘immune-enriched’ vs B ‘immune-depleted’). No statistically significant difference in survival was observed between groups.

