## Supplementary figures and images for "Lymph-node transcriptomics define prognostic immune states in mucosal melanoma and reveal IBA1 as a practical biomarker for improved prognosis"

### Supplementary Figure 1

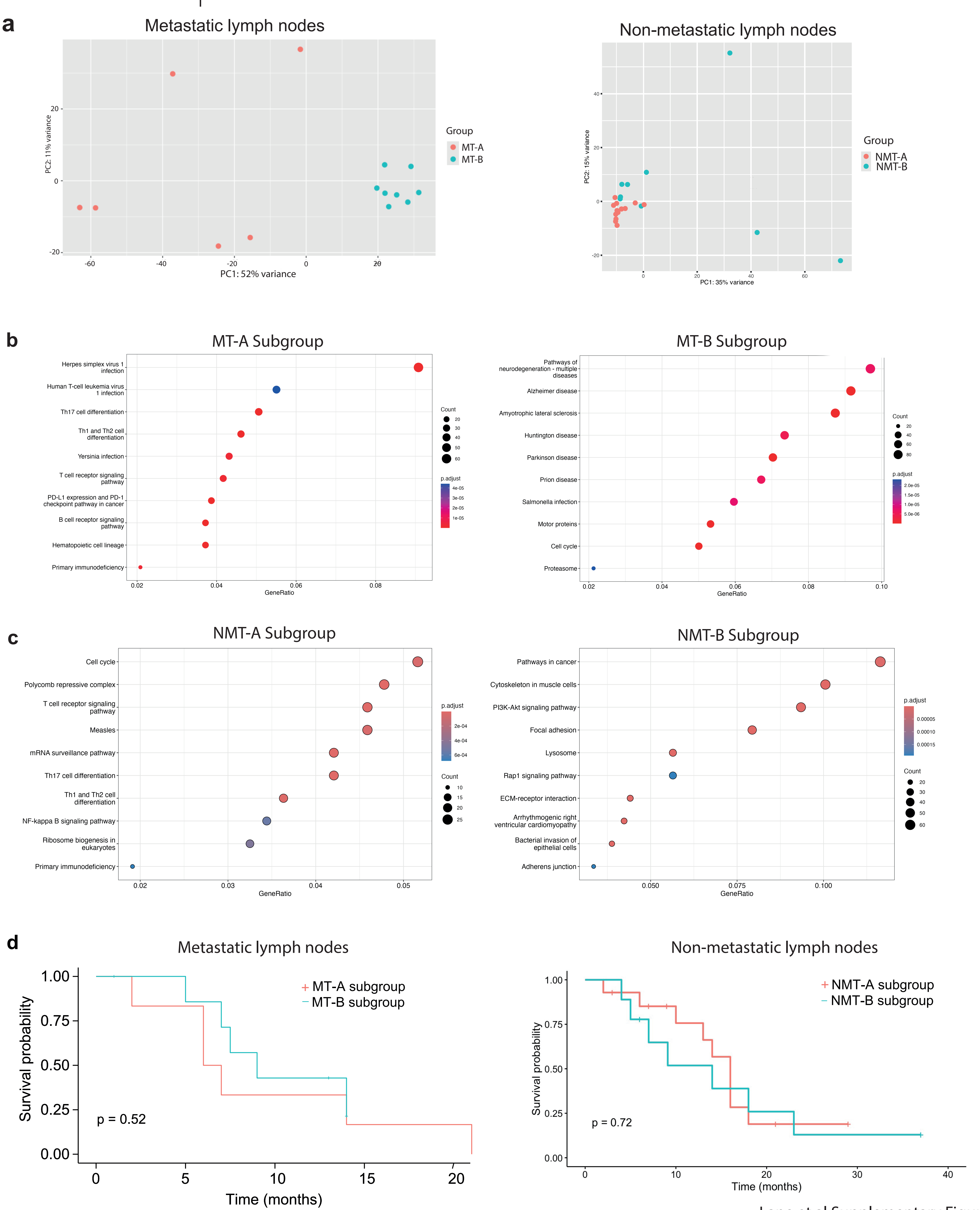


Lane et al Supplementary Figure 1

### Supplementary Figure 2

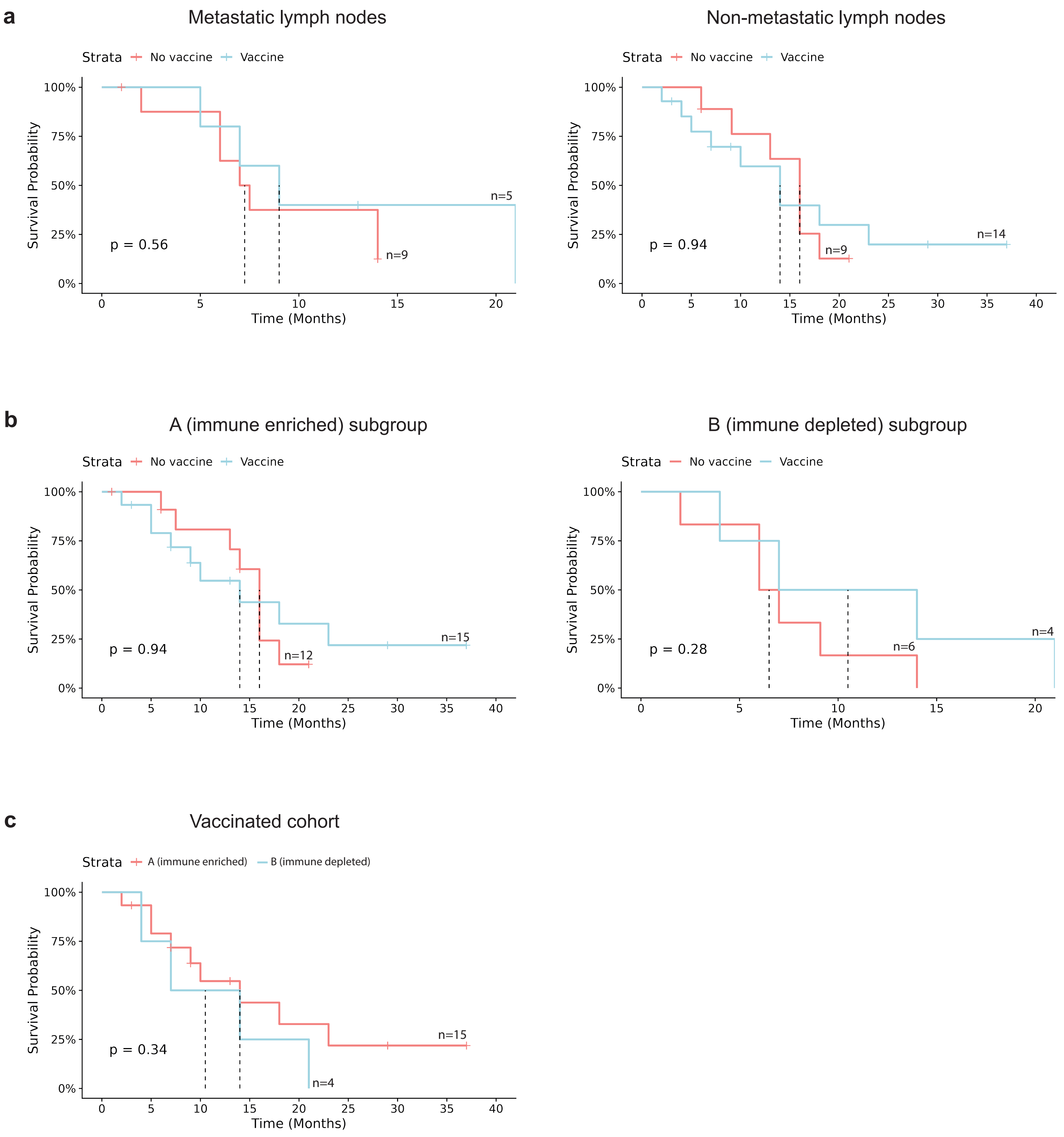


Lane et al Supplementary Figure 2
